# Collateral sensitivities with predictive signatures emerge in a novel model of evolved chemoresistance in osteosarcoma

**DOI:** 10.64898/2026.08.10.743977

**Authors:** Zachary Burke, Kristi Lin-Rahardja, Gabriel Mandel, Jarrell Imamura, Arda Durmaz, Elizabeth Nowak, Masahiro Hitomi, Jacob Scott

## Abstract

Osteosarcoma (OS) is the most common primary malignant bone tumor in children and adolescents. Despite aggressive multimodal therapy, including surgery and chemotherapy with methotrexate, doxorubicin, and cisplatin (MAP), outcomes for patients with relapsed or refractory disease remain poor, with no standardized second-line regimen. We developed a clinically calibrated, temporally resolved *in vitro* model to examine how resistance and collateral drug responses evolve during treatment with methotrexate, doxorubicin, and cisplatin (MAP).

We modeled the evolution of chemotherapy resistance *in vitro* by exposing an OS cell line, MG63.3, to clinically relevant cycles of MAP therapy. MAP resistance and collateral drug responses were quantified over time across five independent evolutionary replicates, and three solvent-treated replicates served as controls. We complemented this phenotypic screening with transcriptomic profiling to extract predictive biomarkers of collateral drug response that could be used to personalize second-line treatment in patients with refractory OS.

MAP exposure produced progressive resistance to doxorubicin and methotrexate, whereas cisplatin sensitivity remained comparatively stable. Repeated temporal screening against 12 additional drugs and combinations revealed three broad patterns of collateral response: progressive resistance, progressive sensitivity, and non-linear or stochastic change. Etoposide resistance emerged consistently, while sensitivity developed toward palifosfamide–etoposide and gemcitabine–docetaxel. Transcriptomic profiling showed dynamic, replicate-specific evolutionary trajectories rather than a single uniform resistance state. To demonstrate one approach for how our large, paired dataset could be used to translate these findings into potentially actionable clinical tools, we extracted predictive gene expression signatures of collateral drug response.

By applying a dynamic, clinically inspired model of chemotherapy resistance in OS, we generated a detailed temporal mapping of collateral drug responses that highlights potential therapeutic windows and can be used inform selection of second-line agents. This approach and the resulting dataset can ultimately support personalized medicine strategies for patients with relapsed or refractory OS.

## Introduction

Osteosarcoma (OS) is the most common primary malignant bone tumor in children and adolescents. Standard of care treatment and patient outcomes have changed little in decades.^1,2^ For patients with localized disease, the standard frontline regimen—methotrexate, doxorubicin, and cisplatin (MAP) with complete surgical resection—has yielded survival rates of 60-70%. Patients with poor histological response and relapsed or metastatic disease have a very poor prognosis with long term survival around 20%.^3^ There are no second-line therapies with consistent efficacy in this group.^4,5^ Novel agents including small molecules, monoclonal antibodies, and antibody-drug conjugates have not been effective to date.^6^ New strategies are necessary to improve outcomes in resistant and relapsed disease.

Targeted therapies and traditional biomarker approaches have been ineffective in OS largely due to the extreme heterogeneity inherent in this disease. Evolutionary oncology approaches have been proposed to overcome this heterogeneity hurdle.^7^ We hypothesize that leveraging convergent evolution – convergence of a heterogeneous population to a common phenotype under a shared stressor – and collateral sensitivity – when resistance to one treatment increases sensitivity to another therapeutic agent – can enable new therapeutic strategies in OS. ^8–10^ When paired with predictive biomarkers, this approach holds potential to enable effective personalized drug selection.^9–11^ Several studies have explored convergent evolution and collateral responses in preclinical cancer models, including Ewing sarcoma, but no studies have utilized this method in OS.^8,9,11–15^ In many ways, this approach is *particularly* well-positioned to extract robust signals from heterogeneous tumors such as OS. Whereas Ewing sarcoma is genetically homogeneous with transcriptional heterogeneity, OS is genomically *and* transcriptionally chaotic, making the convergent evolution/collateral sensitivity approach even more valuable for separating useful signals from noisy data.^17,18^

Several *in vitro* models have been developed to interrogate chemotherapy resistance in OS.^19–22^ However, few effectively model standard of care treatment. Many previous models achieve a very high resistance index (RI, measured as EC_50_ of resistant population/EC_50_ of parental population) by using drug concentrations that are higher than maximally achievable plasma concentrations, continuously exposing cells to drug, or by escalating to doses that are not clinically relevant.^3,20^ These models also lack temporal resolution, limiting analyses to parental and maximally resistant populations. Here we present a novel, clinically relevant *in vitro* model of evolved resistance in OS to elucidate and guide personalized evolutionary treatment strategies. We propose that cycled dosing of MAP within the range of maximum observed clinical plasma concentration more closely models resistance mechanisms and evolutionary trajectories seen in patients. When combined with serial collateral response drug screens and temporal transcriptomic analysis, a clinically calibrated, temporal map of collateral response throughout the evolution of MAP resistance emerges. To capture diverse evolutionary pathways in response to MAP stress, 8 independent evolutionary replicates were created from a treatment-naïve population of the OS cell line, MG63.3. Five of these evolutionary replicates were treated with MAP (referred to as experimental replicates) along with three untreated evolutionary replicates passaged in vehicle solvent in parallel (referred to as control replicates). Finally, we sought to lay the translational foundation for this therapeutic approach by identifying potential predictive biomarkers for collateral sensitivities in a panel of 16 drugs or drug combinations commonly used in osteosarcoma.

## Results

### Resistance was induced to methotrexate and doxorubicin, but not cisplatin

We developed resistance to front-line chemotherapy in five of these evolutionary replicates (referred to as experimental replicates) that were treated for a total of 12 cycles (six exposures each to cisplatin/doxorubicin and methotrexate). Three additional evolutionary replicates (referred to as control replicates) were treated with dimethylformamide (DMF) and dimethyl sulfoxide (DMSO) as vehicle controls for the same number of total cycles. Doxorubicin resistance began to develop between cycles three and six, with all experimental replicates showing at least a two-fold increase in EC50 relative to controls at multiple time points. A similar resistance pattern was observed in methotrexate, with emergence of resistance occurring between cycles six and nine. In contrast, we did not observe consistent resistance to cisplatin (**Fig. 2**). This may be due to the on-and-off exposure of cisplatin throughout the MAP regimen, as opposed to the nature of the cell line itself. In separate experiments, we were able to induce cisplatin resistance in MG63.3 cells that were continuously treated with cisplatin over the course of 2 months (**Fig. S1**). The maximum observed EC_50_s of all MAP agents are reported in **Supplemental Table 1**.

**Figure 1.**
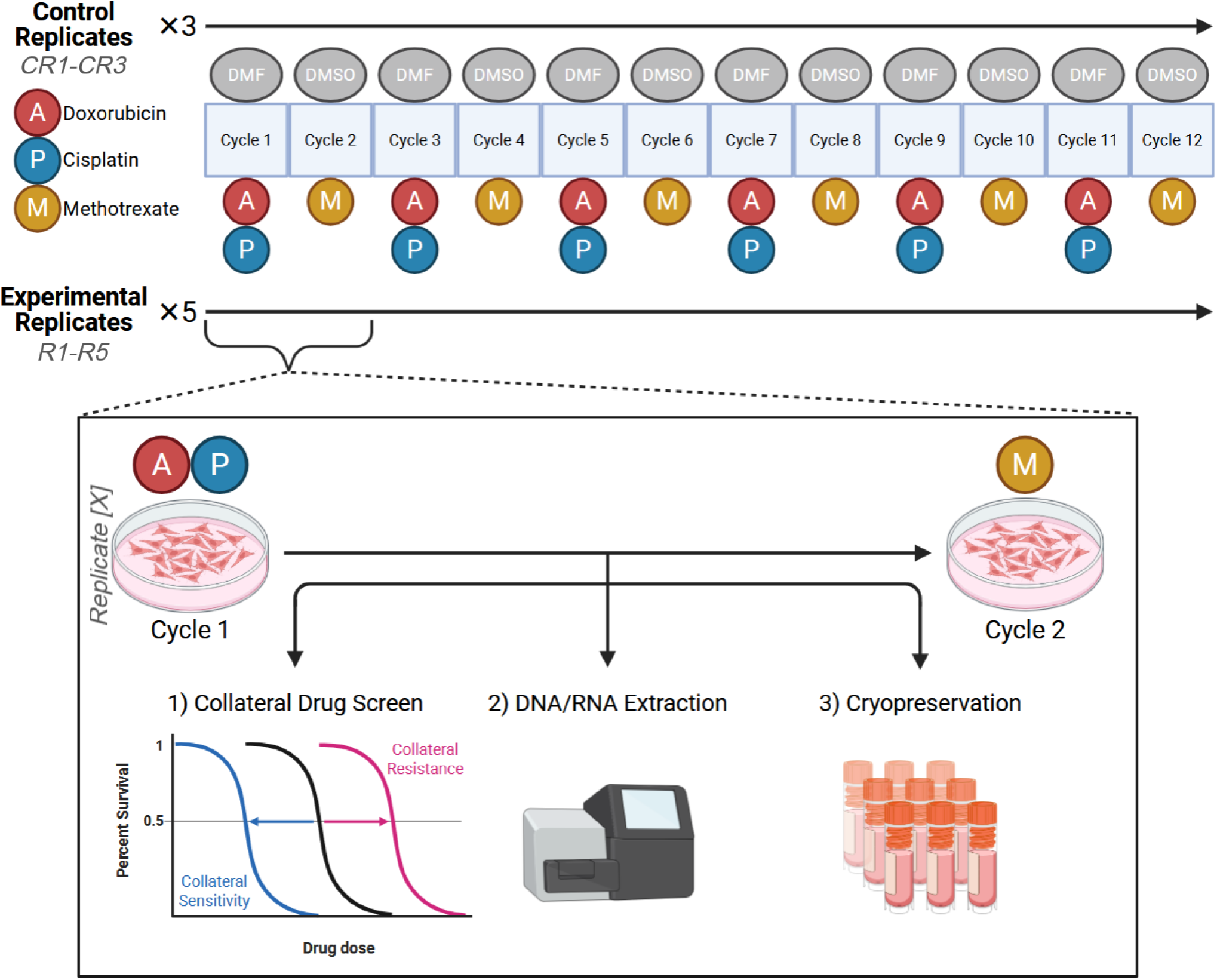
Schema of experimental approach and timeline. Three control replicates (CR1-CR3) are treated with alternating rounds of DMF or DMSO, while five experimental replicates (R1-5) are treated with alternating doses of doxorubicin and cisplatin or methotrexate; each treatment is referred to as one cycle, and all replicates are treated for a total of 12 cycles (six exposures to both cisplatin/doxorubicin and methotrexate). At the end of each cycle, each replicate is screened for collateral drug response, DNA and RNA are extracted, a portion of the replicate population is cryopreserved, and the remaining cells are re-plated to continue onto the next cycle.

**Figure 2.**
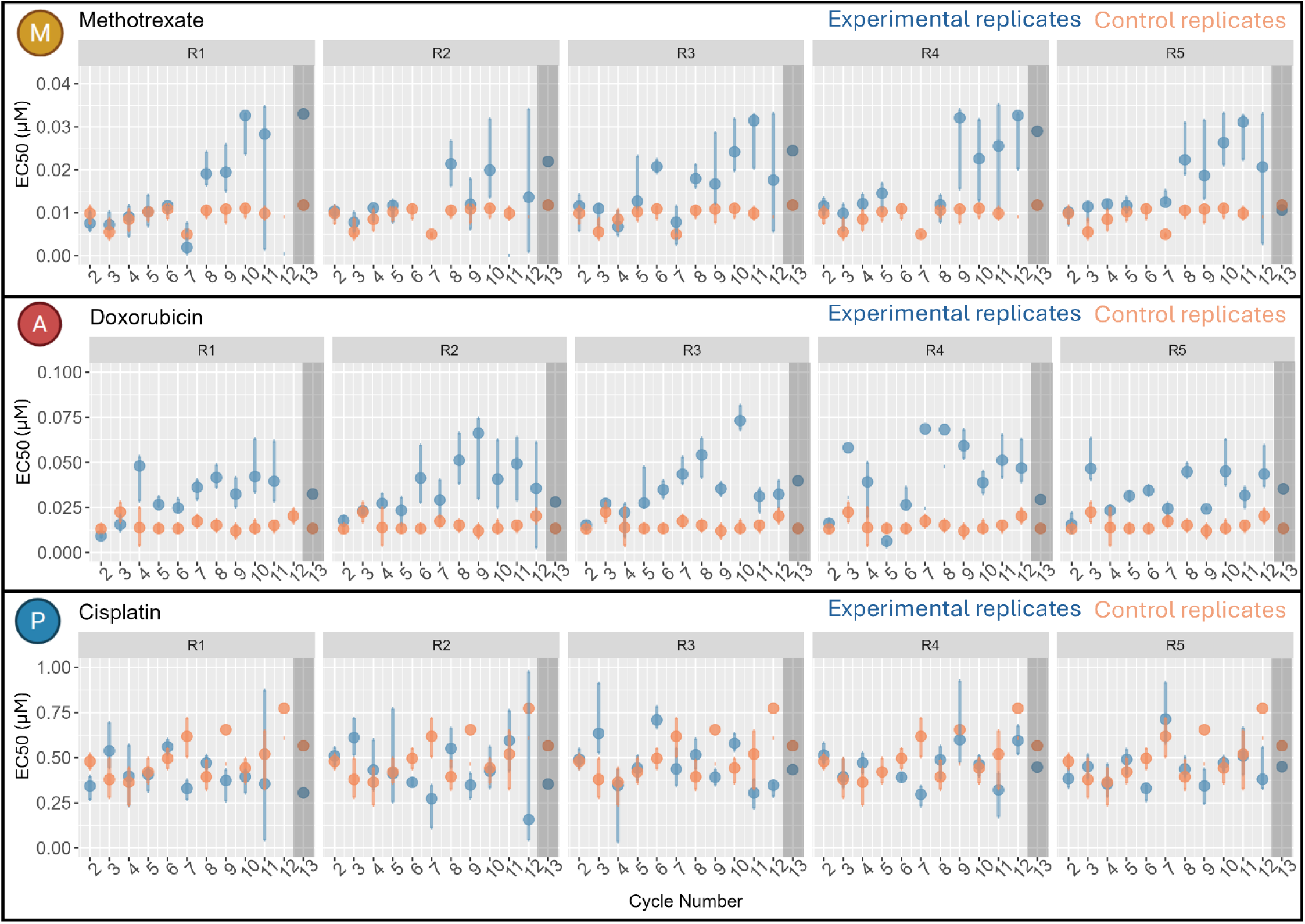
Change in EC_50_ (uM) to the primary drugs (MAP) over time in the five experimental replicates (blue points). Control replicates’ EC_50_s (orange points) are shown as the mean of the three control replicates for that time point. Each row depicts change in drug response to one of the three agents, and each subpanel within a row represents drug response over time in one of the experimental replicates (R1 through R5). All replicates were treated with MAP from cycles 1-12, and cycle 13 (highlighted in gray) included no selection pressure.

To assess whether the resistance phenotypes we observed were stable in the absence of selection, we cultured the cells in drug-free media for an additional passage following the final chemotherapy cycle. The EC₅₀ values for doxorubicin and methotrexate remained higher than controls in all but one replicate (methotrexate R5), confirming persistent chemoresistance after the removal of selection pressure (**Fig. 2**, cycle 13).

### Collateral drug responses are frequent and varied across drugs and experimental time

At all time points for all evolutionary replicates, we measured drug response against twelve drugs or drug combinations commonly used in resistant and recurrent OS. A temporal map of collateral drug responses to each of these treatments is shown in **Fig. 3**.

**Figure 3.**
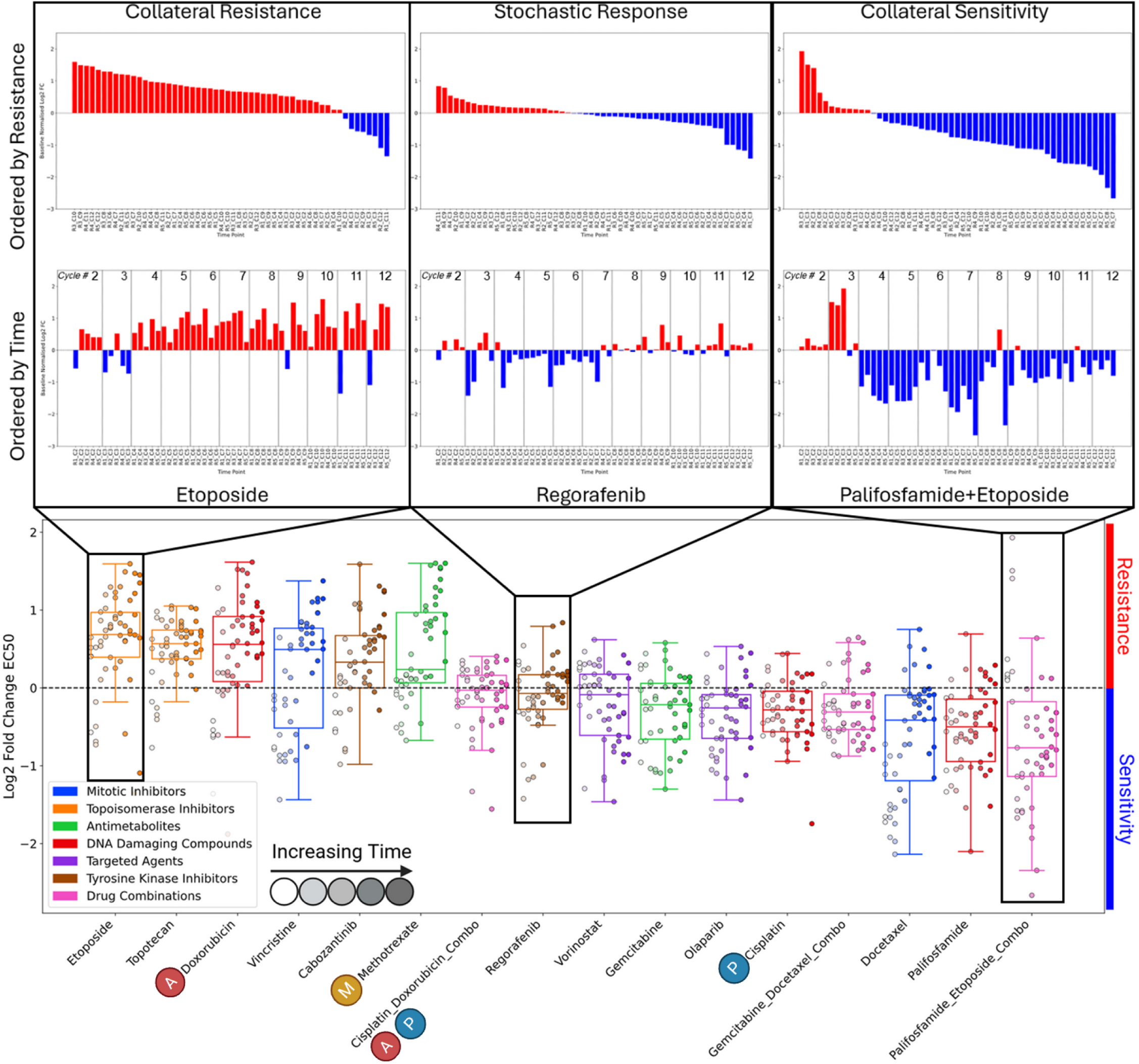
Top panel: Waterfall plots depicting the change in EC_50_ of all replicates (one replicate at one time point per bar) relative to the parental cell line, ordered from most resistant to most sensitive. Three examples are shown of drugs where, from left to right, monotonic collateral resistance emerged, collateral response was temporally stochastic, or monotonic collateral sensitivity emerged. **Middle panel:** Change in collateral response over time across experimental replicates. **Bottom panel:** Box and whisker plot (standard parameters, representing minimum, maximum, quartiles, and median) displaying the log2 fold-change in EC_50_ compared to the parental cell line. Progressively later cycles are indicated by increasing color saturation. Data with control replicates included can be found in **Fig. S2**.

Collateral responses were observed frequently and were varied in magnitude and through experimental time. For all screened drugs or combinations, the experimental replicates had at least one time point with a more than two-fold change in either collateral sensitivity or collateral resistance from the parental line. Only three screened drugs or combinations had replicates that exceeded a four-fold change compared to baseline, all of which were collateral sensitivities (docetaxel, palifosfamide, and palifosfamide/etoposide). Collateral sensitivity was more common than collateral resistance, with eight of twelve screened drugs or combinations showing a median EC_50_ value below baseline. Three general temporal trends were observed across our data: monotonic collateral sensitivity, monotonic collateral resistance, and temporally stochastic responses. For example, we observed monotonic collateral resistance against etoposide and vincristine, which increased as resistance to MAP increased. Monotonic collateral sensitivity developed against the palifosfamide-etoposide combination and vorinostat, with replicates in early time points showing mild collateral resistance, followed by progressive increases toward collateral sensitivity. For many screened drugs and combinations, a temporally stochastic collateral response pattern was observed. For example, replicates showed multiple instances of both collateral resistant and collateral sensitivity against regorafenib that did not progress linearly over time. Interestingly, collateral resistance to etoposide appeared to be mitigated when combined with palifosfamide, as replicates showed a consistent trend toward collateral sensitivity against this combination, in contrast to etoposide alone. A similar phenomenon was seen with cisplatin and doxorubicin combination therapy, where the replicates maintained sensitivity against the combination compared to doxorubicin alone.

### Transcriptomic analyses suggest increased heterogeneity among replicates over time

To assess the temporal dynamics of the molecular changes associated with the evolution of resistance, we performed differential gene expression analysis using an interaction model of time and treatment. We extracted 4 contrasts (control replicates over time, experimental replicates over time, experimental vs. control over time, and average experimental vs. control) and cross-referenced genes of potential interest based on previous literature at FDR < 0.15 level **(Fig. S3)**.^3^ Comparison of the average expression of experimental replicates to control replicates (pooling all the cycles) identified 785 genes with significant differences. Among the cross-referenced genes from previous literature, only *IL32* was found to be differentially expressed in our dataset. We then investigated the temporal expression patterns for genes that have previously been associated with MAP resistance in OS and observed dynamic expression patterns of these genes over time (**Fig. 4**). For instance, ABCB1 and ABCG2 – both drug efflux pumps traditionally thought to *increase* with development or resistance -- showed a strong *decrease* in expression in controls over experimental time, with more moderate changes seen in the experimental replicates. This highlights the non-trivial nature of proper controls in evolutionary studies where the untreated replicates can show transcriptional changes potentially relevant to drug resistance that are difficult to disentangle from the treatment effect. Temporally dynamic expressions are also observed in the evolutionary replicate groups, where *VEGFA*, *BBC3,* and *CXCL8* initially increased in expression from cycles 5 to 10 and then decreased from cycles 11 and 12, highlighting the complexity and temporally dynamic nature of the transcriptional changes.

**Figure 4.**
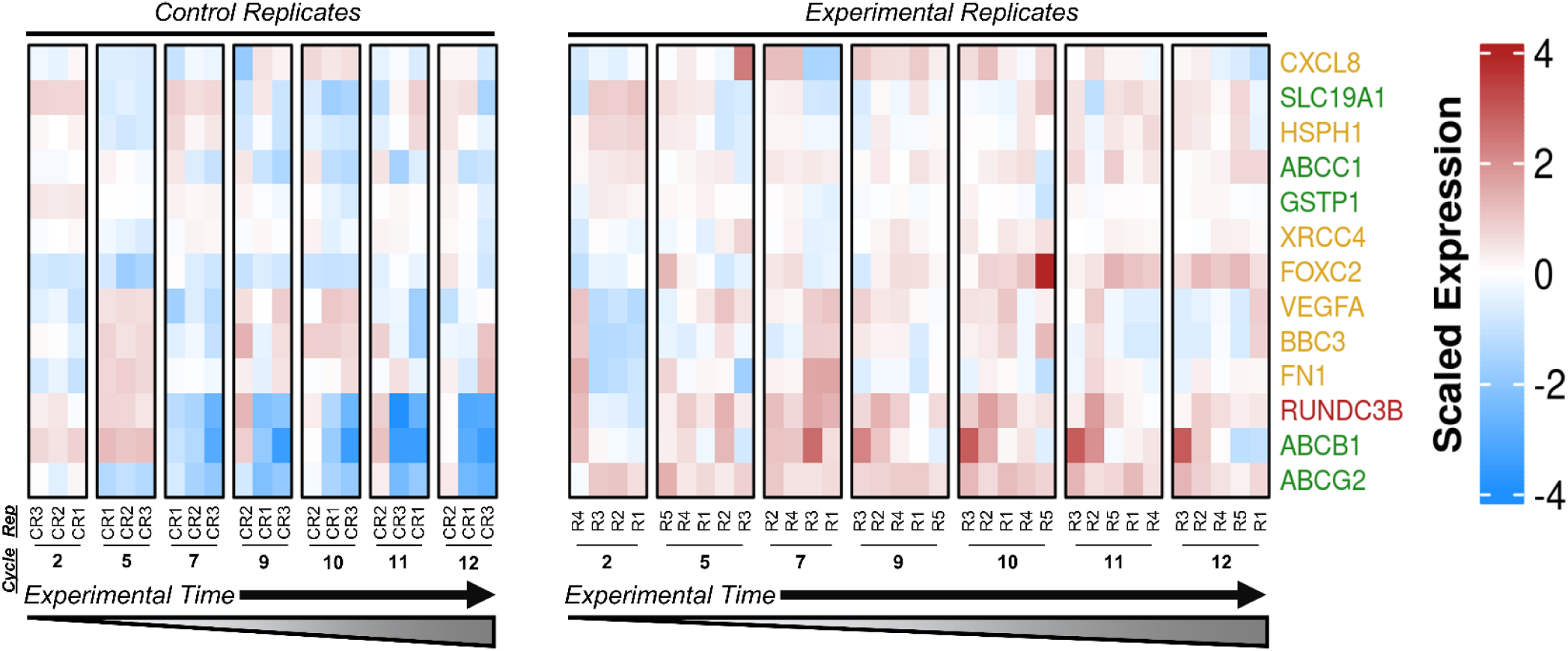
Expression of genes previously associated with MAP resistance in OS. Gene names (rows) are colored based on the significance of comparison (red is significant in both treated and controls, yellow only in experimental replicates, green is only control replicates). Columns are grouped by control replicates on the left, and treated replicates on the right. Columns are subgrouped by cycle number, increasing from left to right. Generally, lower expression of relevant genes was observed in the control replicates compared to the treated replicates.

While the mechanisms of resistance evolution are outside the scope of this paper, the dynamic expression changes of relevant resistance genes over experimental time highlights the importance of temporally resolved models in experimental evolution. To further define the broader transcriptional dynamics, we investigated the global topology of transcriptional and genomic changes through experimental time. We calculated the cosine similarity of the transcriptomes between replicates and found increased heterogeneity across treated replicates toward cycles 9–10, where average pairwise similarities were lower compared to the earlier and later cycles. Some replicates (e.g., R4) that were initially transcriptionally distinct converged by cycle 12, while others (e.g., R1), were initially similar to replicates 2 and 3, diverged into a distinct transcriptional states by cycle 12 (**Fig. 5**). Overall, evolutionary replicates at earlier cycles showed increased similarity to later cycles, suggesting that transcriptional states were not purely linear throughout the emergence of resistance under MAP pressure (**Fig. S5**).

**Figure 5.**
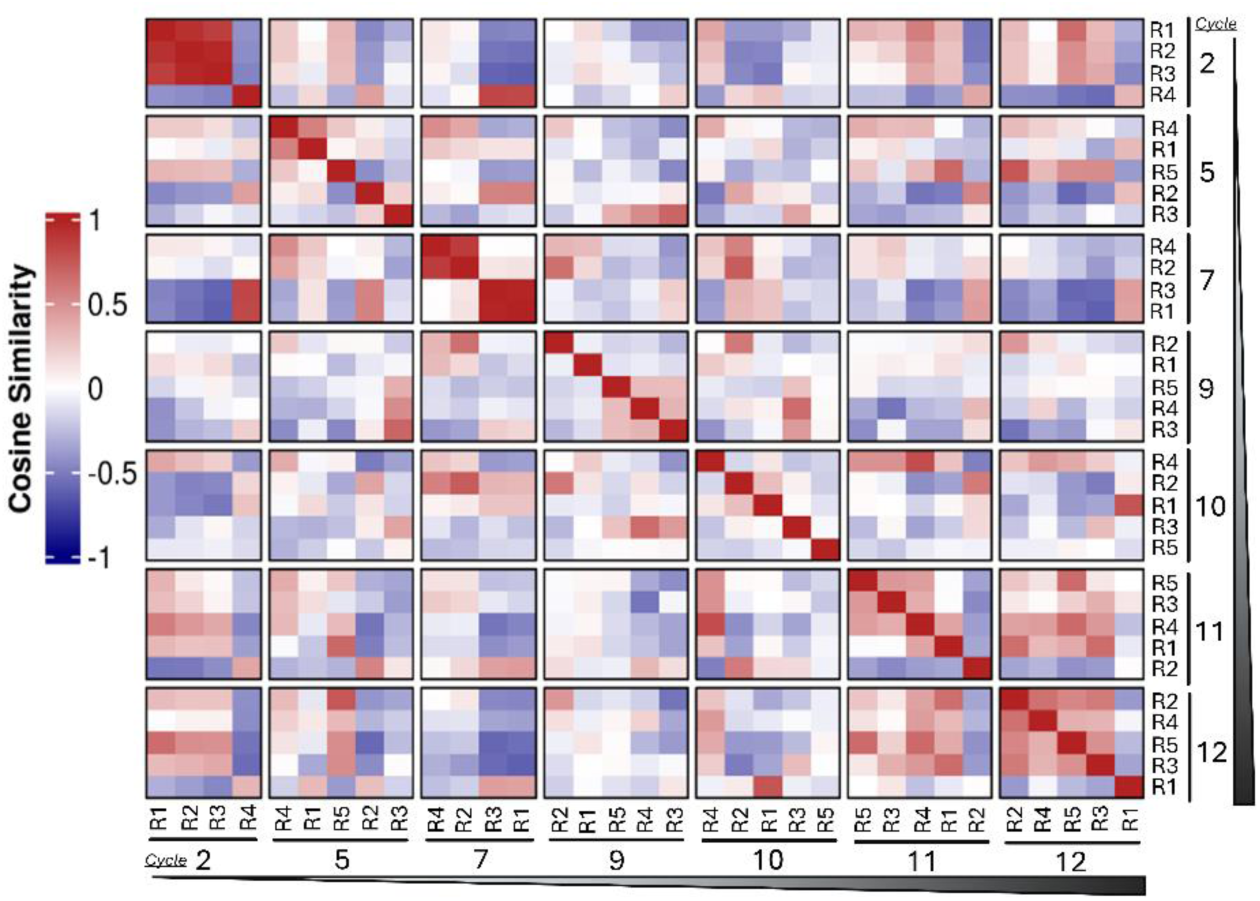
Heatmap illustrating the pairwise cosine similarities of transcriptional profiles for replicates (R1-R5) at different treatment cycles (C2-C12). A cosine similarity approaching 1 indicates high similarity, a value of 0 indicates no similarity, and a value of −1 indicates inverted or opposite similarity. Transcriptional heterogeneity increases towards cycles S-10, indicated by lower average similarity. By cycle 12, some initially distinct replicates converge (e.g., R2), while others diverge to unique states (e.g., R1).

### Gene expression signatures can predict collateral drug response

To extract predictive gene signatures, we adapted the framework demonstrated by Scarborough *et al.*^11^ In short, this convergent evolution-inspired approach used 5-fold cross validation and multiple methods of differential gene expression analysis in 429 epithelial-based cancer cell lines from the Genomics of Drug Sensitivity in Cancer (GDSC) dataset, and filtering through a co-expression network of patient data from the Cancer Genome Atlas (TCGA). These data were used to derive predictive signatures that were robust to cancer subtypes and treatment history. Using our experimental evolution data and transcriptomic data from patient biopsies originating from two independent OS studies in place of the TCGA dataset, we identified 11 signatures (from 16 total screened drugs or combinations) and performed a preliminary assessment of their predictive potential.^23,24^

We first identified differentially expressed genes between any replicates exhibiting clinically relevant collateral response, defined as ≥ 2-fold change in EC_50_ compared to parental. Due to the smaller sample size of our cell line dataset compared to GDSC, we relaxed the stringency of the original method by using a single differential expression analysis method, rather than three. Since gene up-regulation is more statistically robust to detect and is the norm for gene signatures, we retained only the genes that were up-regulated in the collaterally sensitive cohort, and refer to them as seed genes.^11,25,26^ Following this analysis, we retained only the seed genes that ranked within the top 20% of gene co-expression connectivity in both patient datasets.

We applied this approach to all 16 drugs in our screen, which resulted in 11 potential gene signatures (**Table 1**). In four drugs, signatures were unable to be extracted either because there were not enough samples that met our collateral response threshold, or because all genes were ultimately filtered out in the process of extracting the signatures. The signature we extracted for vincristine contained 169 genes, making it impractical for translation -- most published gene signatures with clinical utility have less than 100 genes.^27–32^. Among the 11 candidate signatures we identified, four of these related to drug response against the selection agents in our study: cisplatin, doxorubicin, cisplatin and doxorubicin in combination, and methotrexate. The remaining signatures related to collateral drug response. While the four signatures for the MAP agents are not indicative of collateral drug response by definition, they may still be useful to inform clinicians when MAP therapy is no longer effective and when treatment should be adjusted.

**Table 1.** Candidate translational signatures for collateral response were extracted from 11 of 1c drugs or combinations, including 7 of the 12 secondary drugs screened for collateral response. Underlined/blue drugs are the MAP/selection agents.

| Drug | # Sig Genes |
| --- | --- |
| <u>Methotrexate</u> | 8 |
| <u>Cisplatin</u> | 24 |
| <u>Doxorubicin</u> | 7 |
| <u>Cisplatin+Doxorubicin</u> | 13 |
| Regorafenib | 28 |
| Etoposide | 24 |
| Gemcitabine | 13 |
| Vorinostat | 10 |
| Olaparib | 8 |
| Docetaxel | 4 |
| Palifosfamide+Etoposide | 2 |

To assess the predictive potential of our signatures, we used the signature genes as predictors in a linear regression model to predict EC_50_. To observe each signature-model’s accuracy, we compared the predicted vs. observed EC_50_ against the drug associated with the signature. We also calculated empirical p-values by comparing the *R^2^* of the signature-models to a null distribution, allowing us to compare the performance of our evolution-inspired signatures to 1000 randomly generated gene signatures (each with the same number of genes as the evolution-inspired signature in question).

Through this analysis, we found several signatures with promising preliminary results (**Figure 6** and **Supplemental Table 2**). While this analysis was performed within the dataset from which we extracted the signatures, the extraction process only included the small fraction of samples that exhibited changes in collateral drug response ≥2-fold change EC_50_ in either direction from the parental cell line. For example, 4 out of 50 samples were used to generate the regorafenib model, leaving 46 replicates as the validation set. Across the dataset, the linear regression models fitted with the regorafenib and olaparib signatures were able to predict EC_50_ with notable accuracy and outperformed all but a small fraction of the 1000 random gene signatures’ regression models. This was also the case with the doxorubicin signature, suggesting that this biomarker could be informative of whether first-line chemotherapy should be halted early or extended on a patient-to-patient basis. Ultimately, these results demonstrate that several of our signatures show promise, and with expanded datasets and validation, could become valuable tools for personalizing second-line chemotherapy in patients with resistant or recurrent OS.

**Figure 6.**
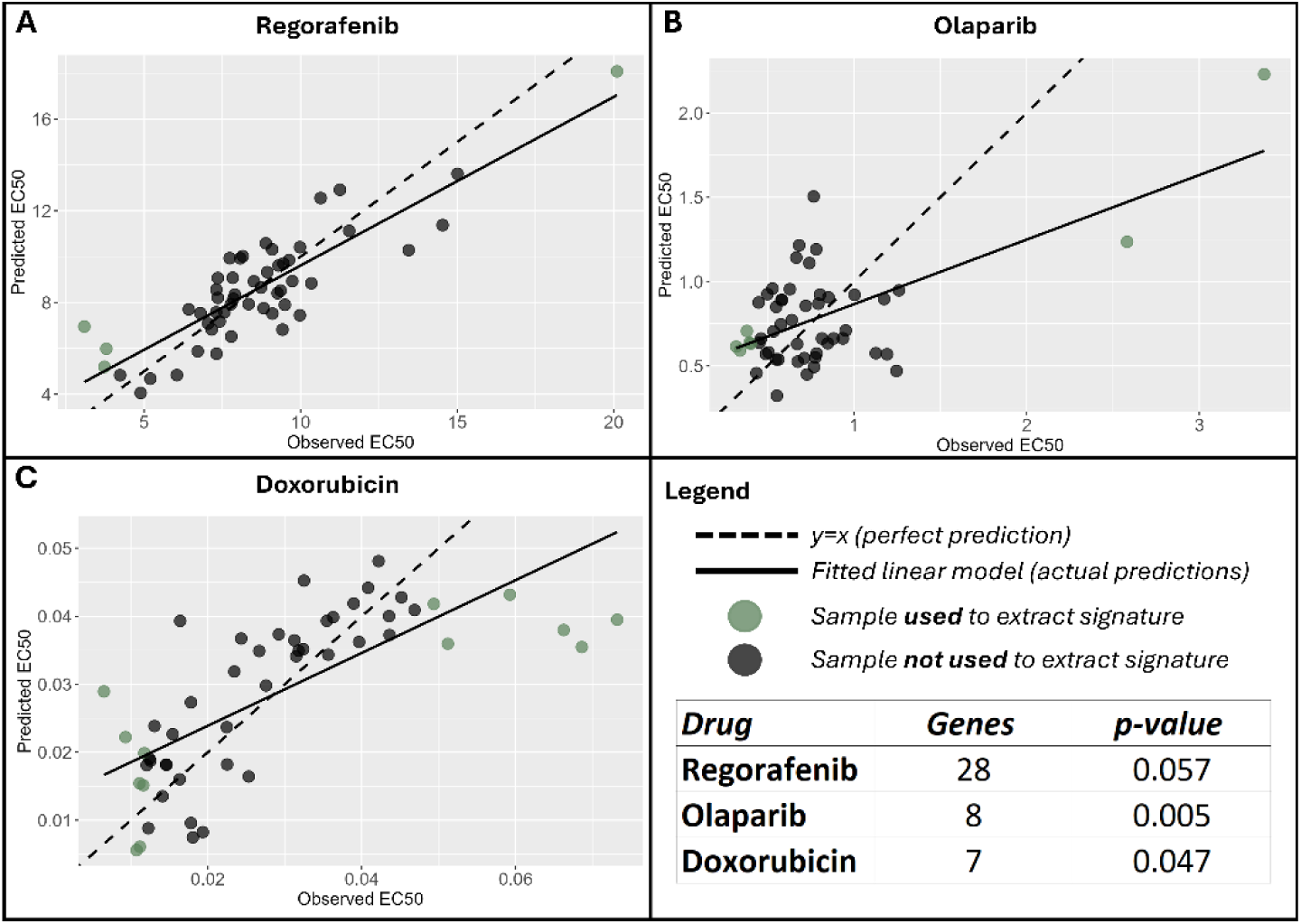
Linear regression<u>s</u> using signature genes to predict EC_50_ demonstrate predictive power of three signatures. All samples were used to fit the linear regression. Samples used for signature extraction are noted in green. **Panels A-C:** show observed versus predicted EC_50_ for regorafenib, olaparib, and doxorubicin respectively. The p-values calculated by comparing the accuracy of signatures to a null distribution of 1000 randomly generated signatures are indicated in a table beneath the legend.

## Discussion

Here we present the first clinically calibrated, temporally resolved map of collateral responses in osteosarcoma. This *in vitro* model shows dynamic collateral responses to a panel of clinically relevant drugs, and leverages convergent states of collateral sensitivity to extract gene expression signatures that predict collateral response. These findings highlight the potential of a convergent evolution/collateral sensitivity approach to canonically heterogeneous malignancies such as OS and lay the foundation for a novel personalized approach to treatment in this disease.

### Collateral drug responses are common and varied in direction and consistency

Of the second line drugs or combinations in our drug panel, two thirds showed showed an overall trend toward collateral sensitivity in our dataset. However, instances of collateral sensitivity were not uniformly durable and were often temporally stochastic. There were no apparent trends in collaterally sensitive drugs based on drug class. The two drugs showing the most collaterally resistant response were etoposide and topotecan, both topoisomerase inhibitors. This pattern suggests that while MAP resistance mechanisms may broadly confer cross-resistance to structurally or mechanistically related agents, collateral sensitivities still emerge to a panel of mechanistically diverse drugs, but are less predictable and more likely to be transient.

Notably, collateral resistance developed against etoposide across all our experimental replicates. This may have been mediated by decreased expression of *TOP2B*, a gene targeted by etoposide. In contrast to this trend observed for etoposide alone, we repeatedly saw collateral sensitivity against the combination of palifosfamide and etoposide. Ifosfamide and etoposide (IE) combination therapy is a frequently used regimen for relapsed OS, and palifosfamide is an *in vitro* analog for ifosfamide. While there have been no randomized trials studying combination IE against single agent ifosfamide, retrospective data supports using high dose ifosfamide (9-15 g/m^2^) as a second line regimen with response rates up to 60%.^33^ The stark difference in collateral responses across our replicates between these two treatments highlights the importance of combination therapy.

Another demonstration of the apparent benefit of drug combinations can be seen between gemcitabine and docetaxel, where the combination of these two agents showed more collateral sensitivity than either agent individually. This phenomenon has also been observed in two other OS cell lines, MG-63 and HOS-143B, and this combination is also often used as a second-line regimen for resistant and recurrent OS.^19,34^

### Sensitivity to cisplatin remained stable

While the experimental replicates developed resistance to doxorubicin and methotrexate, sensitivity to cisplatin remained stable across experimental and control replicates. Other studies on MAP resistance in OS have shown varying results, some showing no resistance to cisplatin, while other demonstrate clear cisplatin resistance patterns.^1, 21^ These differences are most likely due to variations in experimental design, such as the strength of selection (i.e. the dose of MAP agents used to drive resistance) or pre-existing resistance in the cell lines used. In our study, additional experiments showed that a continuous, long term exposure to cisplatin was able to induce resistance. This suggests that the lack of resistance against cisplatin in MG63.3 may have been due to the dosage we used in the experiment as well asinnate resistance (Fig S1). While our evolving experimental replicates showed continued sensitivity to cisplatin in MAP-resistant tumor cells, in a clinical setting, patients cannot receive more cisplatin due to cumulative nephrotoxicity, ototoxicity, and neurotoxicity.^35,36^ This raises the question if other, less toxic, platinum agents (carboplatin, oxaliplatin) may continue to be effective in drug resistant OS. Future studies using this model may include examining the possible synergy of cisplatin and doxorubicin in the drug-naive cells and experimental replicates.

### Transcriptomic changes were temporally dynamic and heterogeneous across replicates

Serial transcriptomic profiling across evolutionary replicates suggested that resistance to MAP was not characterized by a single, linear expression trajectory. Genes previously associated with OS chemoresistance exhibited transient and, in some cases, temporally dynamic changes across treatment cycles. For example, expression of *VEGFA*, *BBC3*, and *CXCL8* generally increased during the middle treatment cycles before decreasing at later time points. Other resistance-associated genes, including *ABCB1* and *ABCG2*, also changed over experimental time, but these changes were larger and only significant in the control group, emphasizing the importance of a control in evolutionary experimentation.

The global transcriptomic analysis similarly showed that the five experimental replicates did not follow a uniform evolutionary path. Pairwise transcriptomic similarity decreased near cycles 9–10, indicating greater heterogeneity among independently evolving populations during this period. By cycle 12, some replicates became more transcriptionally similar despite having divergent states earlier in the experiment, while another replicate diverged into a comparatively distinct state at the end of the experiment. A shared phenotype across MAP-treated replicates thus did not require convergence on a single, stable transcriptional state. Instead, multiple transcriptional trajectories appeared to support adaptation under the same selection pressure.

Broadly, the dynamic transcriptional trajectories observed here are consistent with the variability seen in collateral drug responses over time, although the analyses presented in this study do not establish a causal relationship between particular expression states and drug sensitivity.

### Predictive biomarkers of collateral drug response varied in potential translatability and predictive potential

We show that while the genomic and transcriptomic heterogeneity inherent in OS presents formidable challenges for personalized therapy strategies, the specific strength of a convergent-evolution, collateral-sensitivity approach is in extracting robust signals from noise. The collateral sensitivity signatures presented here are not only a promising translational strategy, but serve as a proof-of-concept for the potential benefit of employing evolutionary approaches in OS.

Among the 7 collateral drug response signatures we extracted, the biomarkers for olaparib and regorafenib showed the most promise based on a preliminary assessment within our dataset. Previous studies have shown regorafenib to be a potentially useful agent for patients with refractory OS: two clinical trials showed that regorafenib could benefit a substantial portion of patients with metastatic recurrent OS, but not all. ^37–39^ Additionally, more than half the patients in both trials who received the drug experienced severe adverse events related to the treatment. No clincal trials have evaluated olaparib alone in osteosarcoma, but a current COG-NCI phase II trial evaluating the efficacy of olaparib in combination with ceralasertib is underway. With further refinement and validation, our predictive gene signatures of collateral sensitivity could enable clinicians to distinguish patients with MAP resistant osteosarcoma who are likely to benefit from those who would not, thereby reducing unnecessary exposure to potentially severe adverse events.

Gene signatures may not be necessary to predict collateral drug response in all cases— in cases of consistent collateral resistance or sensitivity, where there is a clear, repeated trend across evolutionary replicates towards either collateral resistance or sensitivity, further testing may not be needed to decide whether that drug would be useful. For instance, monotonic collateral resistance to etoposide emerged in all treated replicates and increased as MAP therapy progressed. If this trend is also observed consistently in patients, additional testing would not be necessary to determine whether etoposide monotherapy should be avoided. A predictive signature would be the most informative in temporally stochastic responses, where collateral drug response fluctuates non-linearly over time. In our experiment, response to olaparib, gemcitibine, and regorafenib was varied among replicates, and response oscillated between collaterally resistant and collaterally sensitive. If this variability is also present in the patient population, a signature predictive of collateral sensitivity to these drugs would be a valuable tool for personalizing therapy after MAP resistance.

### Limitations

This experiment was conducted using a single OS cell line, which may limit the generalizability of our findings and robustness of the collateral drug response biomarkers we extracted. However, parallel assessment of multiple evolutionary replicates allows the capture of diverse evolutionary trajectories, even within one cell line. In the future, our experimental method could be applied to other OS cell lines or patient-derived tumor biopsies, and the resulting data could be compared with our findings in this study. This would strengthen the translational potential of our conclusions, particularly the robustness of the predictive biomarkers we extracted. To this end, we are currently recapitulating our evolution experiment at a smaller scale using several low passage patient derived cell lines.

An additional limitation was the detection of mycoplasma contamination after cycle 7 of chemotherapy exposure across all replicates. Although the contamination was promptly addressed and eradicated using Plasmocin and subsequent testing confirmed clearance, we cannot exclude the possibility that transient infection may have influenced gene expression profiles, cellular physiology, or drug sensitivity during the affected cycles.

## Conclusion

We developed and characterized a clinically relevant and temporally resolved *in vitro* model of chemotherapy resistance in OS, revealing distinct trajectories of drug response and transcriptional dynamics. Through temporal collateral screening, we identified both monotonic and stochastic collateral responses to common second-line agents, including frequent collateral sensitivities. These findings support the potential of using dynamic collateral response profiling to guide second-line therapy selection. Importantly, several of our evolution-inspired predictive gene signatures show promise with preliminary assessments, and with further investigation, may be useful for personalizing second line treatment in patients with MAP resistant disease. Ultimately, our experimental framework provides a foundation for future efforts to personalize treatment in OS, or any other highly heterogeneous malignancy, based on the evolution of treatment resistance and collateral sensitivity profiles.

## Materials and Methods

### Cell Culture

MG63.3 cells, derived by Ren *et al*., were generously gifted by Berkley Gryder and the cell line identity was validated by short tandem repeat testing.^40^ Patient-derived cells were collected from patients who consented to our IRB-approved institutional biobank (Case Western Reserve University IRB #24-812). Cells were maintained in Dulbecco’s Modified Eagle Medium (DMEM) supplemented with 10% Fetal Bovine Serum (FBS) and 1% penicillin and streptomycin at 37°C under humidified atmosphere containing 5% CO_2_.

### Drugs

Cisplatin, doxorubicin, docetaxel, etoposide, gemcitabine, methotrexate, olaparib, palifosfamide, and topotecan were purchased from Cayman Chemical. Cabozantanib, regorafenib, vincristine, and vorinostat were purchased from MedChemExpress. Drugs were dissolved in dimethyl sulfoxide (DMSO), aside for cisplatin and doxorubicin which were dissolved in dimethylformamide (DMF).

We chose to use palifosfamide (Cayman Chemical) (isophosphoramide mustard-lysine tris, ZIO-201) rather than ifosfamide or cyclophoshamide since it represents the active alkylating moiety —ensuring uniform, reproducible exposure in vitro without the need for bioactivation or introduction of off-target metabolites.^41^

For drug response testing, a concentrated drug stock solution was prepared at various concentrations based on optimized dose response curves and drug solubility (**Supplemental Table 3**). Drug combinations were designed based on individual dose–response curves. For each pair—cisplatin with doxorubicin, gemcitabine with docetaxel, and palifosfamide with etoposide—drugs were combined at a fixed 1:1 ratio of their respective EC₅₀ values. Compounds were serially diluted threefold across nine rows of a deep well 96 well plate (ThermoScientific, cat. 260251), with the final well containing only the compound’s respective solvent as a negative control. These dilutions were aliquoted into 96 well master plates (Greiner, item no. 655087) with 8 drugs or drug combinations per plate and stored at –20 °C.

### *In vitro* combination drug treatments to induce drug resistance

MG63.3 cells were plated at 250,000 cells per 15 cm diameter dishes (Costar, ref. 353003) per evolutionary replicate. A total of eight evolutionary replicates were created. Five experimental replicates were exposed to the standard-of-care drug cycles for five days, washed with phosphate-buffered saline (PBS), and subsequently re-incubated with growth media without drug. After recovery to sub-confluency at the end of each cycle, the replicates were trypsinized and partitioned for downstream applications, including cryopreservation for archival storage, DNA/RNA extraction, drug sensitivity screening, and reseeding for subsequent chemotherapy cycles. While clinically the last two doses of doxorubicin are not accompanied by cisplatin, a protocol deviation was made in our experiment to continue cell exposure to cisplatin to attempt to induce resistance.

Treatment was initially administered at the EC₁₀ concentrations of cisplatin (0.07315 uM) and doxorubicin (0.02993 uM) and the EC₅₀ concentration of methotrexate (0.1705 uM). Doses were started at this lower cis/dox EC50 due to combination therapy and complete eradication of cells with higher dosing. After six cycles of chemotherapy, in the absence of significant observed resistance to methotrexate or cisplatin, the doses were escalated to the EC₃₀ concentrations for cisplatin (0.2042 uM) and doxorubicin (0.02528 uM), and the methotrexate dose was doubled (0.341 uM). All doses used in this experiment were lower than previously observed maximum serum concentrations for these agents.^42^

The three control evolutionary replicates were treated with matching concentrations (by volume) of vehicle solvent (DMF or DMSO) used for the experimental replicates. To ensure that controls experienced comparable culture conditions, they were passaged during the recovery periods of the experimental replicates and the downstream applications were performed at corresponding time points. Although exact synchronization was not always possible, efforts were made to align control and experimental replicates by evolutionary time, meaning both sets of cultures underwent the equivalent number of drug or drug vehicle exposure. This design helped account for potential batch effects arising from prolonged culture or handling variability.

### Dose Response Assays

Growth assays were performed to determine optimal seeding density for a 6-day assay in 96-well plates (Costar, ref. 353003), which was established as 600 cells per well. On day 0, cells were seeded at a density of 600 cells in 90 μL of culture medium per well. On Day 1, master drug plates were thawed at room temperature, gently agitated to ensure homogeneity, and diluted 1:100 (or 1:33 for palifosfamide, olaparib, and vorinostat) in culture medium, depending on the drug. After agitation, a volume of 10 μL of each drug dilution (resulting in a final dilution factor of 1:1000 relative to the original stock, or 1:333 for palifosfamide, olaparib, and vorinostat) was added in triplicate to the plate inoculated with cells on day 0. Cells were incubated for an additional five days. On Day 6, 11 μL of Alamar Blue reagent (Thermo Fisher) was added to each well and incubated for 2 hours. Fluorescence was measured using a Tecan plate reader with excitation at 560 nm and emission at 590 nm. Cell viability background-corrected using Alamar Blue-treated wells with media only/no cells, then normalized to the average signal from triplicate wells treated with the corresponding vehicle control (DMF or DMSO) for each drug. Dose response assays for cycle 1 were lost due to contamination.

### Dose Response Analysis

To determine the net Alamar Blue signal, the mean background fluorescence of the triplicated wells with no cell inoculation from each plate was subtracted from the raw fluorescence of each well. These corrected values were then normalized to the average signal of the vehicle controls to calculate the survival fraction S(x). Each drug was then fitted to a four-parameter log-logistic (LL.4) model of the form:

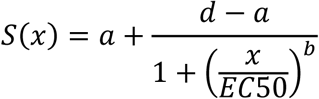

where *S(x)* represents the modeled survival fraction at dose *x*; d and a denote the upper and lower asymptotes, corresponding to the minimum and maximum response levels as x approaches 0 and ∞, respectively. EC_50_ is the inflection point of the concentration at which the response is halfway between *a* and *d*; and *b* is the Hill slope, which governs the steepness of the curve at the inflection point. Under this parameterization of the LL.4 model, a positive *b* yields a monotonically decreasing response curve, while a negative *b* produces a monotonically increasing profile. Within each biological replicate, bootstrapping was performed to generate 500 resampled datasets. The resulting distribution was used to estimate the median EC_50_ and corresponding 95% confidence interval for that replicate.

### Mycoplasma Testing and Treatment

Testing was performed regularly using Lonza kit as per manufacturers’ instructions. Mycoplasma contamination was identified after cycle 7 of treatment across all 8 replicates and was eradicated using Plasmocin (InvivoGen) as per manufacturer’s instructions prior to moving forward in experiment. Subsequent screenings for mycoplasma were negative.

### DNA and RNA extraction

DNA and RNA were extracted from cells using AllPrep DNA and RNA extraction kit as per manufacturers protocol (Qiagen). RNA was sequenced by the Cleveland Clinic Genomics Core. DNA after chemotherapy cycles 2, 5, 7, 9, 11, and 12 were sent to Novogene for sequencing.

### Transcriptome Quantification

Transcriptomic profiling was performed using STAR-Salmon framework with the Human Genome Build 38 (hg38) analysis set as the reference genome. Raw reads were first trimmed using *fastp* and then aligned to hg38 analysis set reference using STAR aligner in two-pass mode. Subsequently, Salmon was used for quantification whereas the *tximport* package was used to aggregate transcript-level transcripts per million (TPMs) to gene-level length bias removed counts.

### Transcription factor activity Inference

We estimated transcription factor (TF) activity using the TIGER (Transcriptional Inference using Gene Expression and Regulatory data) package.^43^ The CollecTRI collection is used as the network encoding TF-target interactions, with parameters signed and *tfExpressed* set to true.

### Differential expression analysis

Differential gene expression across samples was performed using *edgeR* package.^44^ We used natural-cubic splines with a single knot set at cycle 5 timepoint via *splines* package to model linear and non-linear expression changes across different cycles of MAP treatment. The effect of treatment is modeled as an interaction term between the spline basis vectors and treatment status.

### Signature Extraction

Signature extraction was performed in R. One signature was extracted for each drug included in the collateral drug screen. Drug response data was used to identify what samples showed at least a 2-fold increase or decrease in collateral drug response, and these samples were then compared with differential expression analysis. Genes found to be differentially up-regulated in sensitive samples were kept as “seed genes”.

Co-expression was calculated in two independent clinical datasets. The data from Hu *et al.* included RNAseq for 70 osteosarcoma patients of various ages and tumor stages.^23^ The data from Buddingh *et al.* included microarray expression for 34 patients of various ages whose tumors had eventually metastasized.^24^ Co-expression was determined within each dataset by calculating Spearman correlation between each pairwise combination of all genes across all samples in the dataset. The Spearman correlation was then converted to percentiles and binarized (1 if the pairwise-correlation was in the top 5% of co-expressed genes, 0 if it was not).

To filter the seed genes to maintain genes with high co-expression in patients, we subsetted the binarized matrix so rows contained seed genes, and columns contained all genes. For each gene (column), we then calculated the proportion of seed genes that were highly co-expressed with it to calculate a connectivity score (ex. For gene A, if 4/10 seed genes were in the top 5% of co-expression, the connectivity score for gene A is 0.4). Once a connectivity score was calculated for all genes (columns) in the dataset, we identified genes with a connectivity score in the top 20%. Finally, we identified which seed genes were present in this top 20%. These identified genes serve as the final signature.

### Linear Regression Modeling

Linear regression models were fitted using the *lm* function from the *stats* package in R.^45^ For each of the 11 signatures, one model was fit using the z-score normalized expression values of signature genes as predictors for EC_50._ The *predict* function from the *stats* package was used to apply the fitted model to the data and generate predicted EC_50_ for all samples. To summarize the predictive accuracy of the signature model, the *R^2^* of predicted vs. measured EC_50_ was recorded.

Per signature, a one-sided empirical p-value was calculated by generating a null distribution comparing the *R^2^* of the signature model to *R^2^* values of linear regression models fitted with 1000 unique randomly generated gene signatures of the same length. The empirical p-value was calculated using the equation below, where *r* represents the number of random signature models with an *R^2^* above the signature model, and *n* represents the number of random iterations (1000).

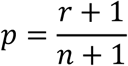

## Author Contributions

Conceptualization: GM, ZB, JS

Methodology: GM, BN, ZB, AD, MH

Experimentation: GM, BN, JI, MH

Analysis: AD, JI, KL

Manuscript: ZB, GM, AD, KL, JI, JS

## NIH T32

### NCCN

This work was funded by the NCCN Foundation®. Any opinions, findings, and conclusions expressed in this material are those of the author(s) and do not necessarily reflect those of National Comprehensive Cancer Network® (NCCN®) or the NCCN Foundation® Cleveland Clinic Velosano Grant

## Supplemental Material

**Supplemental Figure 1:**
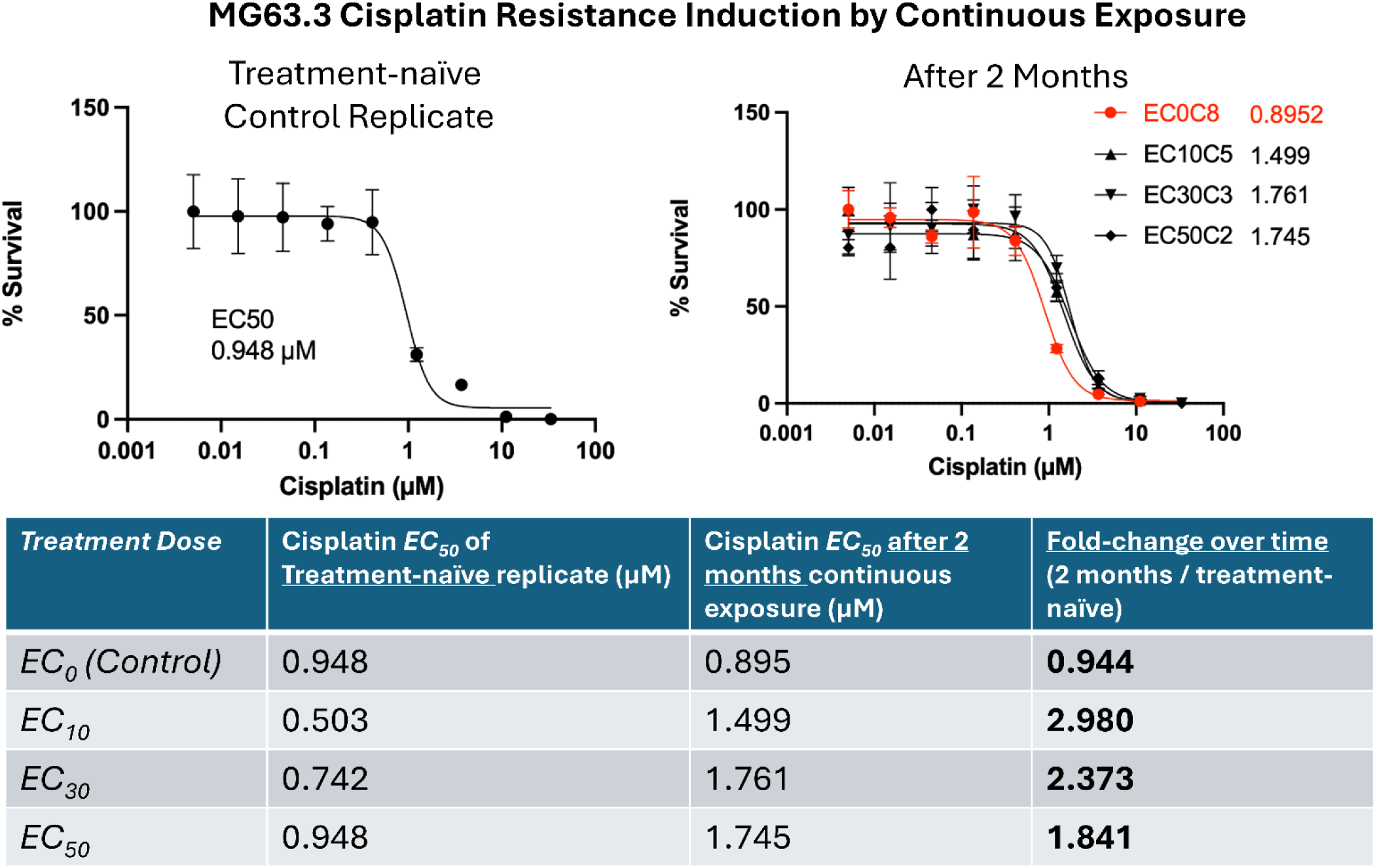
Cisplatin resistance was inducible in MG63.3 via continuous exposure. Four replicates (independent from the evolution experiment presented in this study) were treated with four different doses of cisplatin for 2 months, and the cisplatin dose responses before and after this period were measured. Roughly 2-fold increases in cisplatin EC_50_ were observed in the treated replicates compared to the untreated control, and little difference was observed among the three treated replicates.

**Supplemental Figure 2:**
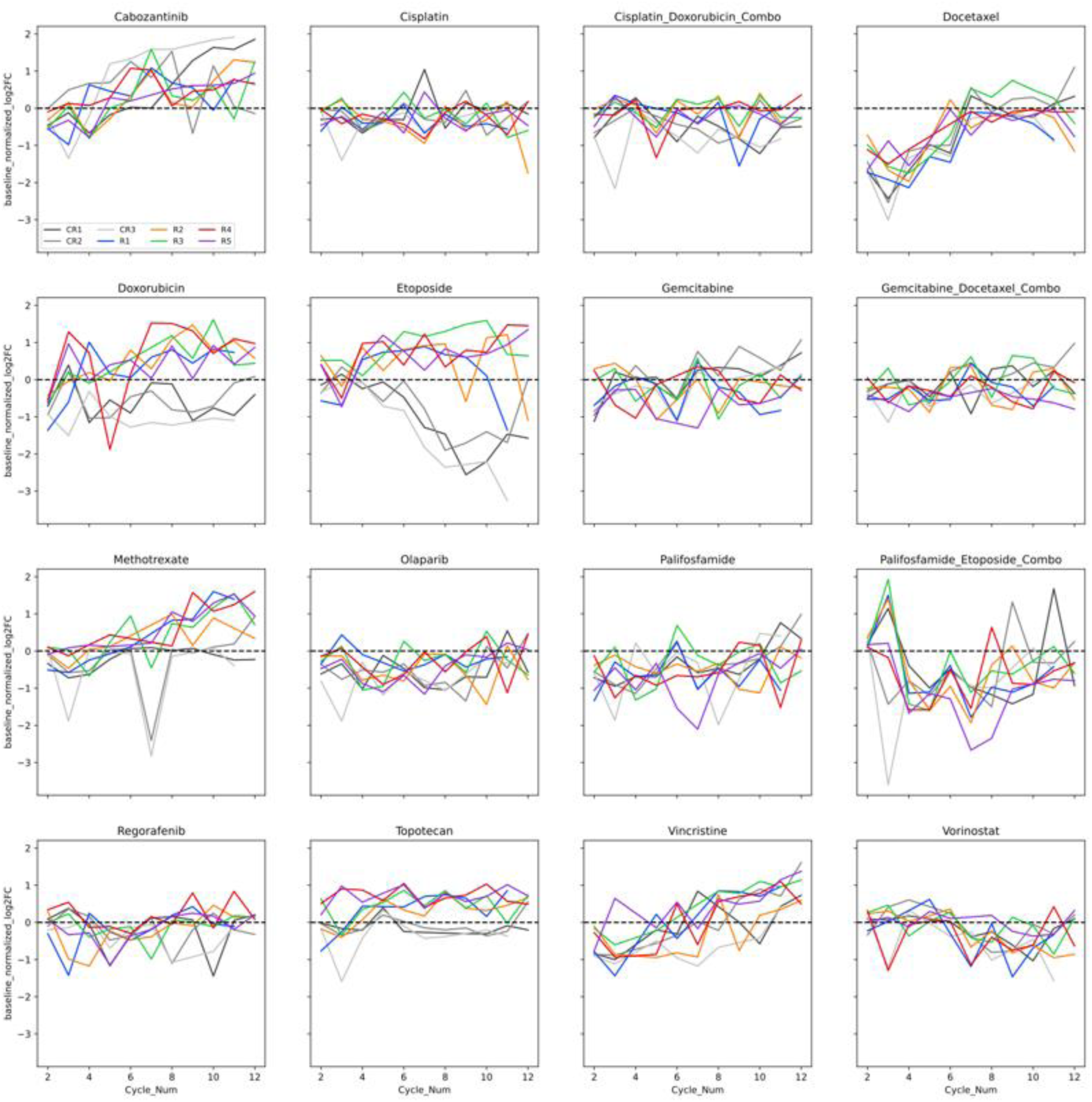
Log2(fold-change in EC_50_) compared to parental across drugs and combinations, including control replicates. Colored lines represent experimental replicates, while grayscale lines represent control replicates.

**Supplementary Figure 3.**
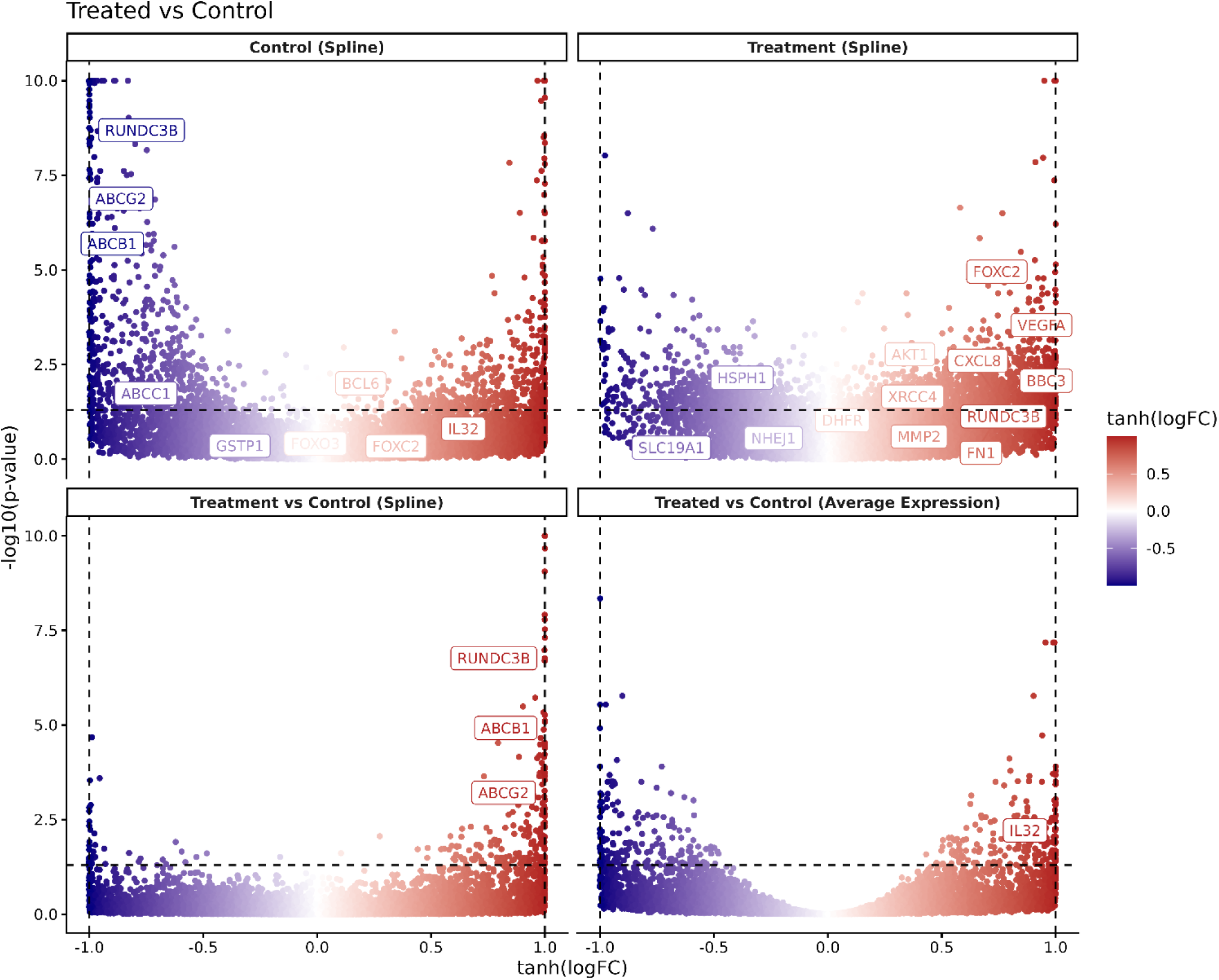
Volcano plots and corresponding pathway enrichment results for the 4 comparisons showing both global differences and changes associated with cycle progressions. Comparisons tagged with “spline” are showing the linear/non-linear associations of expression over time, whereas the comparison tagged with “average expression” is showing the global differences pooling samples from all the cycles together. The genes labeled are genes of potential interest associated with MAP resistance based on previous studies and with significant difference in the corresponding comparison at q-value < 0.15 significance level.

**Supplementary Figure 4.**
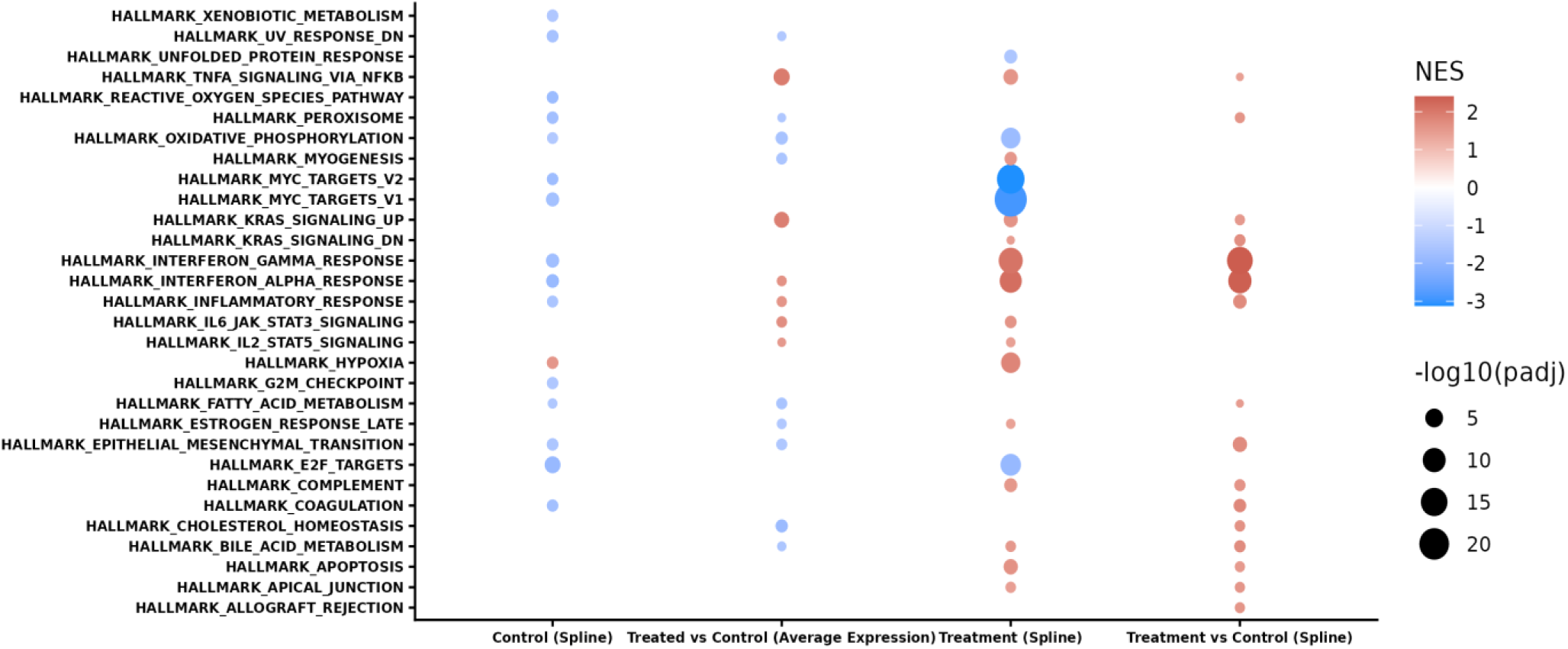
Gene expression enrichment scores comparing averaged expression of control replicates and treated replicates (labeled as “Average Expression”) or comparing control replicates and treated replicates over time (labeled as “Spline”). Color indicates increased enrichment (red) or decreased enrichment (blue), and point size indicates statistical significance.

**Supplemental Figure 5.**
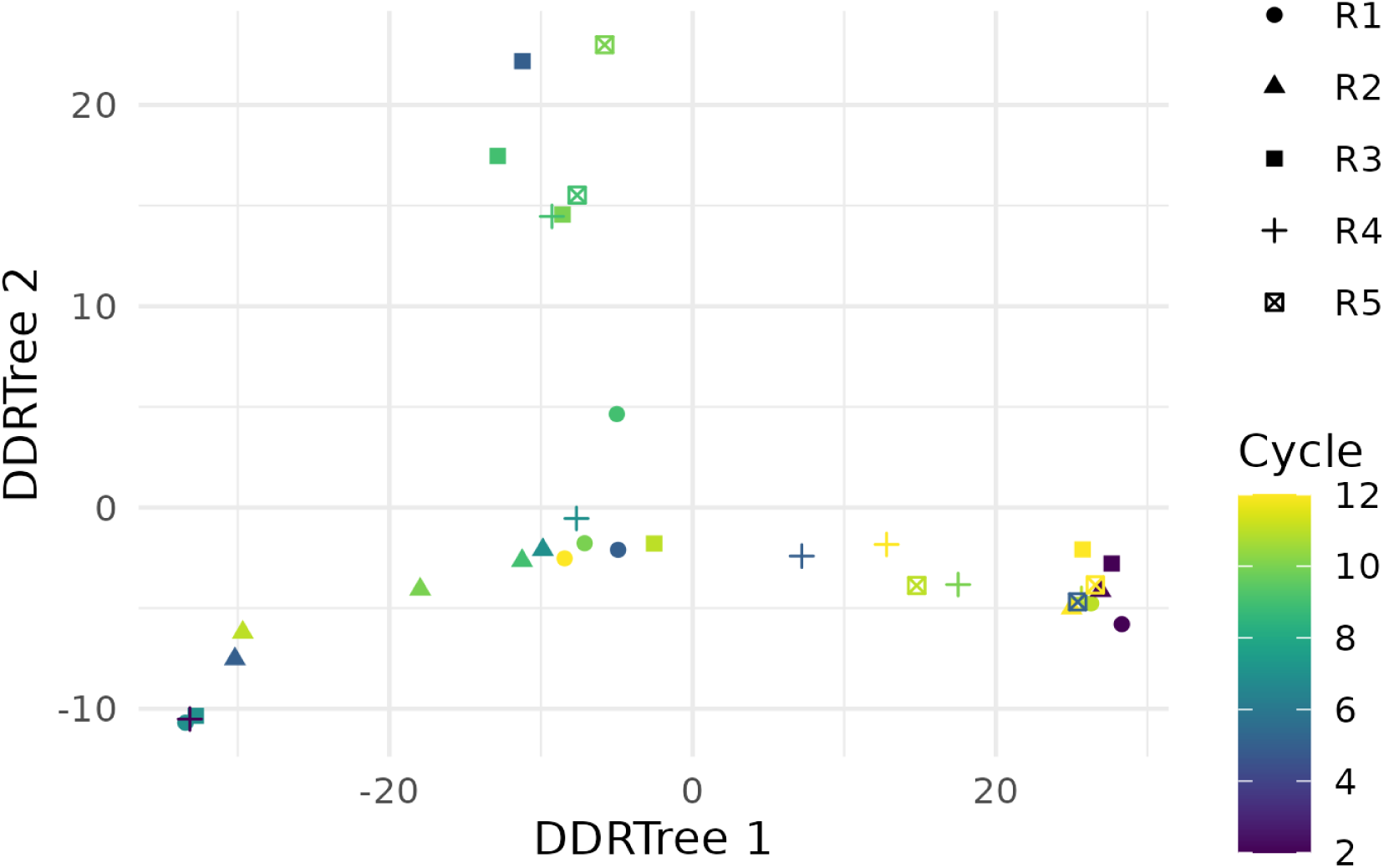
2D principal graph representation of the transcriptional profiles estimated using *DDRTree* which shows higher similarities between Cycle 2 and Cycle 12 replicates indicating higher transcriptional diversity at midpoint cycles.

**Supplemental Figure 6.**
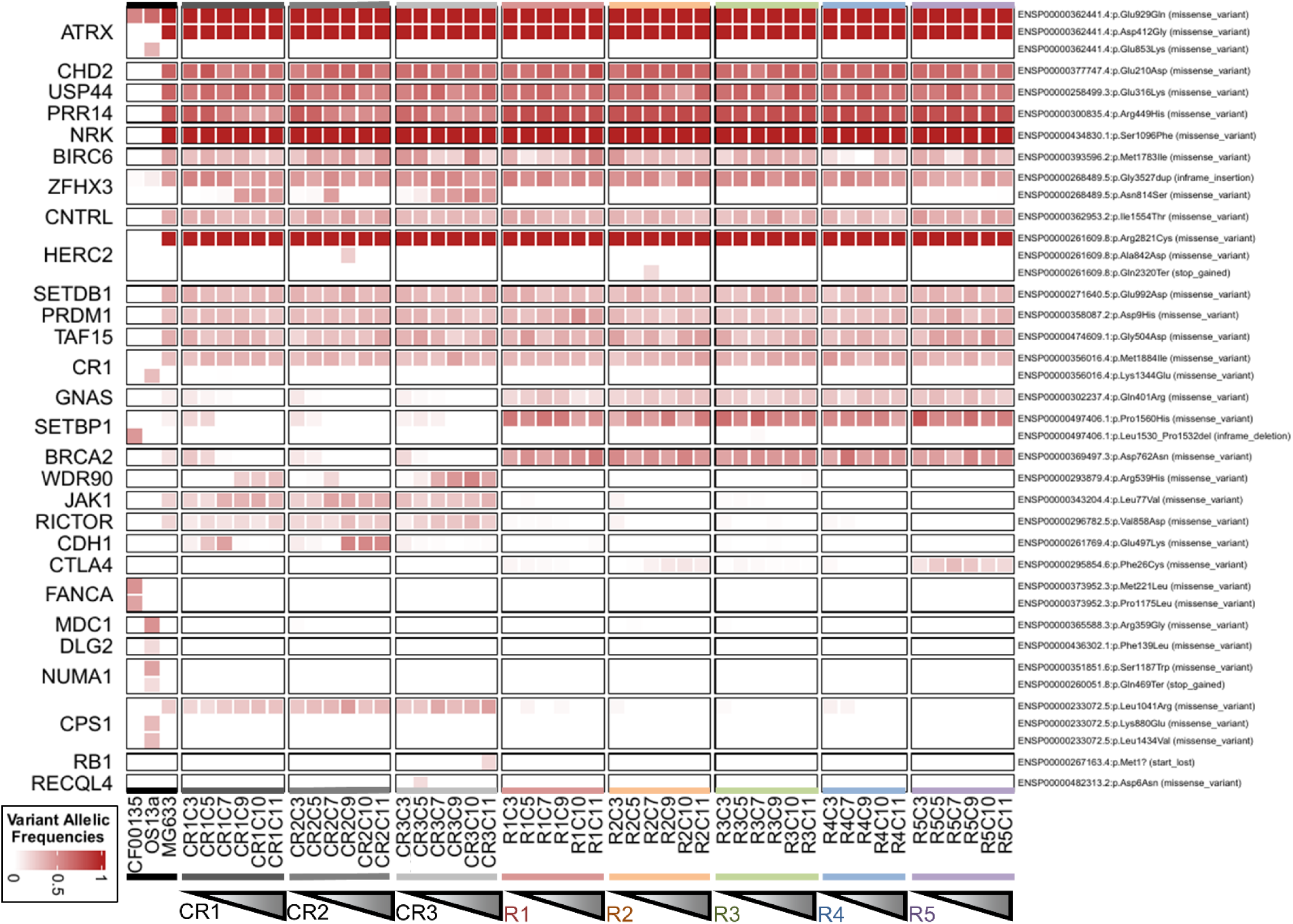
Heatmap of variant allelic frequencies across replicates and time. Replicates are grouped in columns and ordered over experimental time within each group. The first 3 columns represent a tumor biopsy of OS (CCF00135), a separate OS cell line (OS13a), and the parental cell line used in our experiment (MG63.3). Genes with a variant allelic frequency greater than 0.05 are shown by row.

---

**Supplemental Table 1.** Maximum observed EC_50_ of MAP agents for each replicate and what cycle the value was observed at.

| Drug | Replicate | Baseline EC<sub>50</sub><br>(uM) | Max EC<sub>50</sub><br>(uM) | Resistance<br>Index (RI) | Time of<br>Max EC<sub>50</sub> |
| --- | --- | --- | --- | --- | --- |
| Methotrexate | R1 | 0.011 | 1.358 | 126.24 | C12 |
| Cisplatin+Doxorubicin | R1 | 0.158 | 0.768 | 4.87 | C12 |
| Cisplatin | R1 | 0.525 | 3.173 | 6.04 | C12 |
| Doxorubicin | R1 | 0.024 | 0.130 | 5.43 | C12 |
| Methotrexate | R2 | 0.011 | 9.670 | 898.73 | C11 |
| Cisplatin+Doxorubicin | R2 | 0.158 | 0.209 | 1.32 | C10 |
| Cisplatin | R2 | 0.525 | 0.611 | 1.16 | C3 |
| Doxorubicin | R2 | 0.024 | 0.066 | 2.77 | C9 |
| Methotrexate | R3 | 0.011 | 0.031 | 2.92 | C11 |
| Cisplatin+Doxorubicin | R3 | 0.158 | 0.199 | 1.26 | C10 |
| Cisplatin | R3 | 0.525 | 0.708 | 1.35 | C6 |
| Doxorubicin | R3 | 0.024 | 0.073 | 3.06 | C10 |
| Methotrexate | R4 | 0.011 | 0.162 | 15.03 | C6 |
| Cisplatin+Doxorubicin | R4 | 0.158 | 0.202 | 1.28 | C12 |
| Cisplatin | R4 | 0.525 | 3.515 | 6.70 | C5 |
| Doxorubicin | R4 | 0.024 | 0.069 | 2.87 | C7 |
| Methotrexate | R5 | 0.011 | 0.133 | 12.41 | C6 |
| Cisplatin+Doxorubicin | R5 | 0.158 | 0.195 | 1.24 | C3 |
| Cisplatin | R5 | 0.525 | 0.713 | 1.36 | C7 |
| Doxorubicin | R5 | 0.024 | 0.047 | 1.95 | C3 |

**Supplemental Table 2.** Linear regression results for all extracted gene signatures. Selection (MAP) agents are listed in blue.

| Drug | # Sig Genes | R<sup>2</sup> | p-value |
| --- | --- | --- | --- |
| Cisplatin | 24 | 0.539 | 0.714 |
| Doxorubicin | 7 | 0.536 | 0.047 |
| Cisplatin+Doxorubicin | 13 | 0.165 | 0.972 |
| Methotrexate | 8 | 0.181 | 0.378 |
| Regorafenib | 28 | 0.735 | 0.057 |
| Gemcitabine | 13 | 0.501 | 0.369 |
| Vorinostat | 10 | 0.389 | 0.098 |
| Olaparib | 8 | 0.383 | 0.005 |
| Etoposide | 24 | 0.708 | 0.694 |
| Docetaxel | 4 | 0.347 | 0.008 |
| Palifosfamide+Etoposide | 2 | 0.169 | 0.023 |

**Supplemental Table 3.**
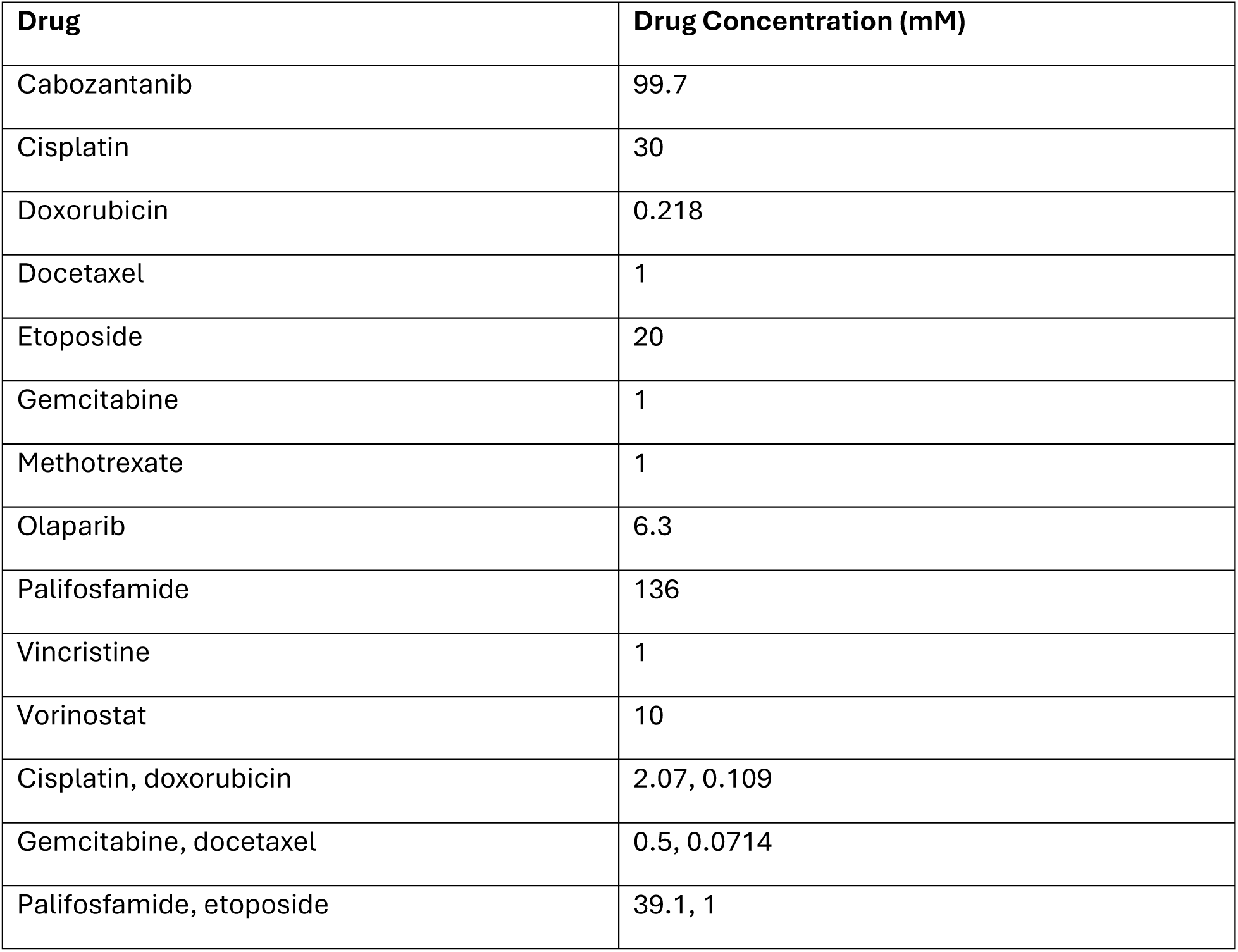
Concentrations of drug stock solutions used in drug response assays.

